# Synaptic turnover promotes efficient learning in bio-realistic spiking neural networks

**DOI:** 10.1101/2023.05.22.541722

**Authors:** Nikos Malakasis, Spyridon Chavlis, Panayiota Poirazi

## Abstract

While artificial machine learning systems achieve superhuman performance in specific tasks such as language processing, image and video recognition, they do so use extremely large datasets and huge amounts of power. On the other hand, the brain remains superior in several cognitively challenging tasks while operating with the energy of a small lightbulb. We use a biologically constrained spiking neural network model to explore how the neural tissue achieves such high efficiency and assess its learning capacity on discrimination tasks. We found that synaptic turnover, a form of structural plasticity, which is the ability of the brain to form and eliminate synapses continuously, increases both the speed and the performance of our network on all tasks tested. Moreover, it allows accurate learning using a smaller number of examples. Importantly, these improvements are most significant under conditions of resource scarcity, such as when the number of trainable parameters is halved and when the task difficulty is increased. Our findings provide new insights into the mechanisms that underlie efficient learning in the brain and can inspire the development of more efficient and flexible machine learning algorithms.

## Introduction

The brain is an incredibly efficient machine as it can process and store vast amounts of information with exceptional speed and accuracy while utilizing the energy of a small light bulb^1,2^. Beyond its energetic efficiency, and unlike most artificial learning systems, the biological brain can also learn continuously over a lifetime^3,4^. It constantly integrates new information with existing knowledge without catastrophically forgetting what was previously learned^5^, forming a complex web of concepts and ideas. This efficient learning, in terms of both energy and performance, is believed to be facilitated by a process known as neural plasticity^6–11^. By forming, eliminating, and fine-tuning connections between neurons^12^, neural plasticity allows the brain to adjust to newly acquired information and affect cognitive functions such as perception^13,14^, motor control^15–17^, and decision-making^18–20^.

Neural plasticity can be divided into two broad and overlapping categories, functional and structural plasticity^21^. Functional plasticity typically refers to changes in the strength of individual synapses in response to neural activity^22^, mediated by alterations in the release probability of neurotransmitters^23^, and the number of receptors on the postsynaptic neuron^24^. Functional plasticity is thought to underlie many forms of learning and memory, including long-term potentiation (LTP) and long-term depression (LTD)^25^. Structural plasticity, on the other hand, typically refers to changes in the overall structure of the nervous system^26^, including the growth of new dendrites, filopodia, and axons^27^, the formation of new and the elimination of existing synapses^28^ as well as changes in the size/shape of spines^29^.

Synaptic turnover, a form of structural plasticity, refers to the dynamic process of continuous formation and dismantling of synapses^30–32^ and is strongly associated with learning in multiple areas of the brain^21,33–38^. Studies have shown that synaptic turnover is vital during development, whereby the formation and elimination of synapses are critical for establishing functional neural circuits^39–41^. However, synaptic turnover is not restricted to developmental periods^36,42^. Several studies have shown that it continues throughout the lifespan^43–45^, providing a mechanism to adjust the wiring diagram of neuronal circuits in addition to the strength of synapses. Similarly, synaptic pruning is important for stabilizing and consolidating memories by removing synapses that are no longer needed^46–48^.

While numerous experimental and modeling studies have investigated the role of neural plasticity in learning and memory, little is known about the role of synaptic turnover, especially when it operates in conjunction with other biological features such as active dendrites and other plasticity forms. Synaptic turnover is especially interesting because of its potential to expand the processing capabilities of sparsely connected networks like those often seen in biological brains. This is achieved by the dynamic formation and dismantling of task-specific microcircuits^49^. Moreover, the ability to rewire through synaptic turnover can dramatically increase the storage capacity in simulated biological networks^50–52^ and enhance the ability to transmit information efficiently across neurons^53^. Experiments in transgenic animals with higher synaptic turnover rates showed improvements in learning a fear conditioning task^35^ and enhanced motor learning ability^54^. Moreover, the higher level of spine turnover is correlated with greater capacity for subsequent song imitation in birds^55^, thus verifying its importance in behaving animals. Computational modeling suggested that the mechanism via which learning was improved resided at the dendritic level, whereby increased turnover led to synapse clustering in active dendrites and sparser memory encoding^56^ and may protect memories from subsequent modifications^57^.

To investigate whether synaptic turnover underlies the learning efficiency of biological circuits, we expanded a biologically constrained spiking neural network (SNN) model^56^ and assessed its learning capacity on various discrimination tasks. We found that the presence of synaptic turnover resulted in higher performance accuracy and faster learning across all scenarios tested. Moreover, these improvements were highest under conditions of resource scarcity, namely when the number of trainable parameters was halved and when the task difficulty was increased. Improvements were due to a more accurate representation of the different image categories by the network weights through their flexible and more efficient utilization. Our results highlight the important role of structural plasticity in optimizing learning in biological circuits and open new avenues for exploring the applicability of such biological plasticity rules in machine learning applications.

## Results

To investigate the impact of synaptic turnover on efficient learning, we modified a biologically constrained computational network of neurons^56,58^ (**Figure 1A**). Our model network consists of excitatory and inhibitory leaky integrate-and-fire neuronal models with adaptation^59^. All neuronal models have independent, nonlinear dendritic compartments modeled as two-stage integrators^58,60^ (**Figure 1B**). The excitatory neuronal population receives external excitation while the inhibitory neurons provide feedback inhibition to the network at both the soma and the dendritic compartments. The inhibition scheme is similar to the soma-targeting and dendrite-targeting interneurons found in the cortex^61,62^. Additionally, the excitatory population is split into two groups, with each group encoding a different class of inputs. Similarly, the inhibitory population is divided into two functional groups: those that regulate network activity via feedback inhibition to both subpopulations (perisomatic and dendritic inhibition) and those that receive excitation from one subpopulation and inhibit the opposite subpopulation (dendritic inhibition). As a result, the winner subpopulation prevents the other from firing, enforcing a Winner-Takes-All mechanism in the network^63^. External input is provided to both subpopulations via spike trains with a mean firing rate analogous to the intensity of each pixel in the sample image (**Figure 1C**). Finally, during the training phase, the two excitatory subpopulations receive class-specific teaching signals that consist of excitatory input to the pyramidal neurons (**Figure 1A**).

**Figure 1.**
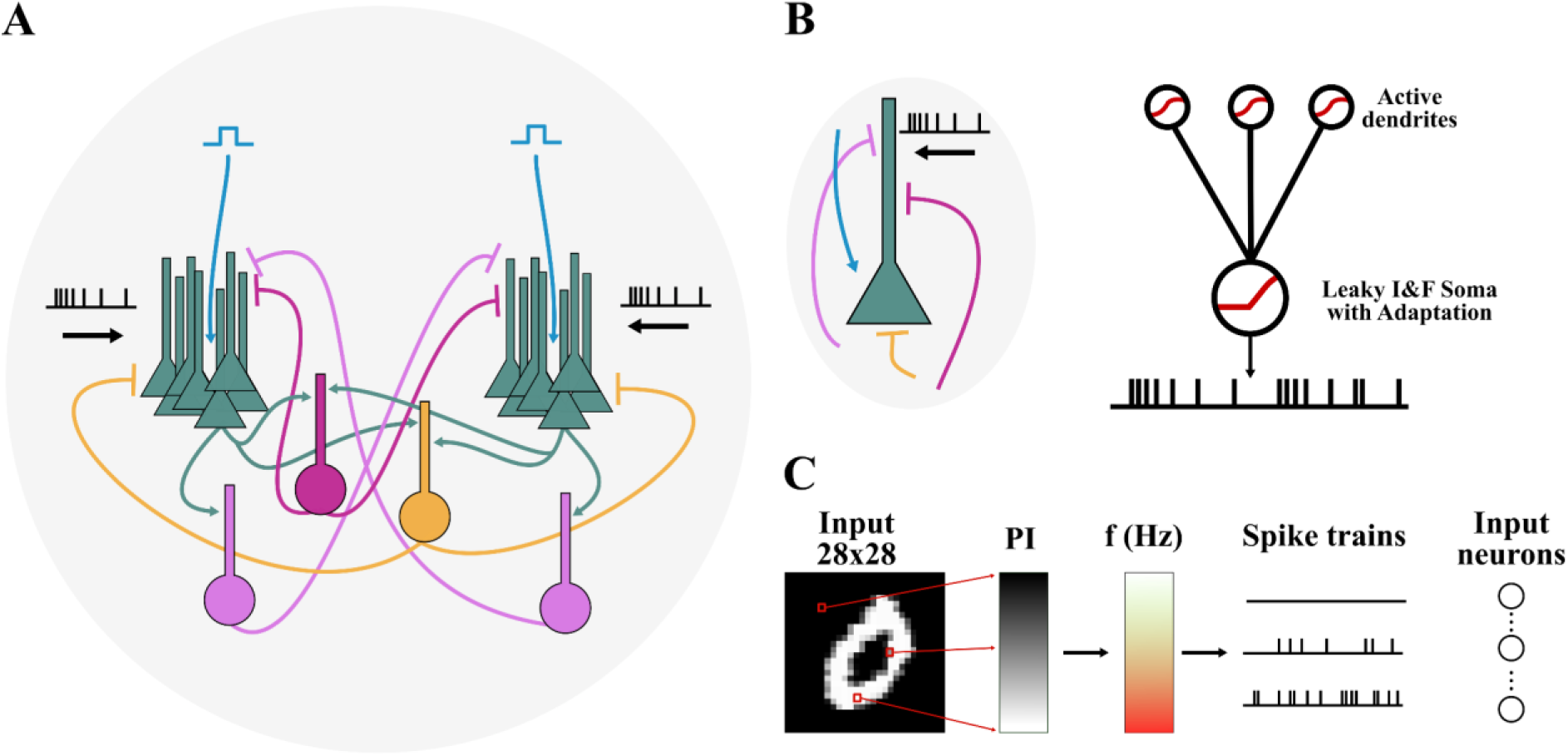
Schematic representation of the spiking neural network. **A.** The SNN model consists of two populations of pyramidal neurons with dendritic compartments (green). Inhibition is provided via somatic targeting (yellow) and dendritic targeting (purple) interneurons. Dendritic targeting interneurons are subdivided into two groups: the first group projects randomly to both subpopulations (dark purple), while the second group projects to the opposite subpopulation (light purple). Each subpopulation receives a teaching signal (light blue) when the input sample matches its target class. **B.** (left) Schematic of a pyramidal model cell and its incoming connections. (right) Each pyramidal cell consists of a somatic compartment and ten dendritic compartments, all of which are furnished with nonlinear activation functions (red). The somatic compartment is simulated as an Integrate-and-Fire model with adaptation, while the dendritic compartment as a nonlinear integrator. **C.** Each input image pixel is translated into spike trains with a mean average firing rate positively correlated with its intensity in a way that pixels with high intensity correspond to spike trains with a high mean firing rate.

The network contains two primary forms of plasticity: functional and structural. Functional plasticity directly affects the strength of the synapses (**Figure 2A**), while structural plasticity operates over longer time scales and leads to axonal re-organization, such that weak synapses are pruned, and new ones are generated onto new postsynaptic sites^36^ (**Figure 2B**). Functional plasticity is in line with the synaptic tagging and capture (STC) model, which requires both the tagging and availability of plasticity-related proteins (PRPs) to strengthen and weaken synapses^64–66^. Calcium levels determine the sign of the synaptic tag (positive or negative) inspired by the Bienenstock-Cooper-Munro (BCM) rule^67^, which then leads to potentiation or long-term depression. Inspired by synaptic plasticity in the brain, we implemented a synapse-specific learning rate in which small synapses change fast while large synapses are resistant to change^68–71^, leading to their stabilization^72^ (**Figure 2D**). As a consequence of stabilization, training stops when the learning rate of at least half of the synapses falls below a specific threshold (see Methods, **Figure 2C**).

**Figure 2.**
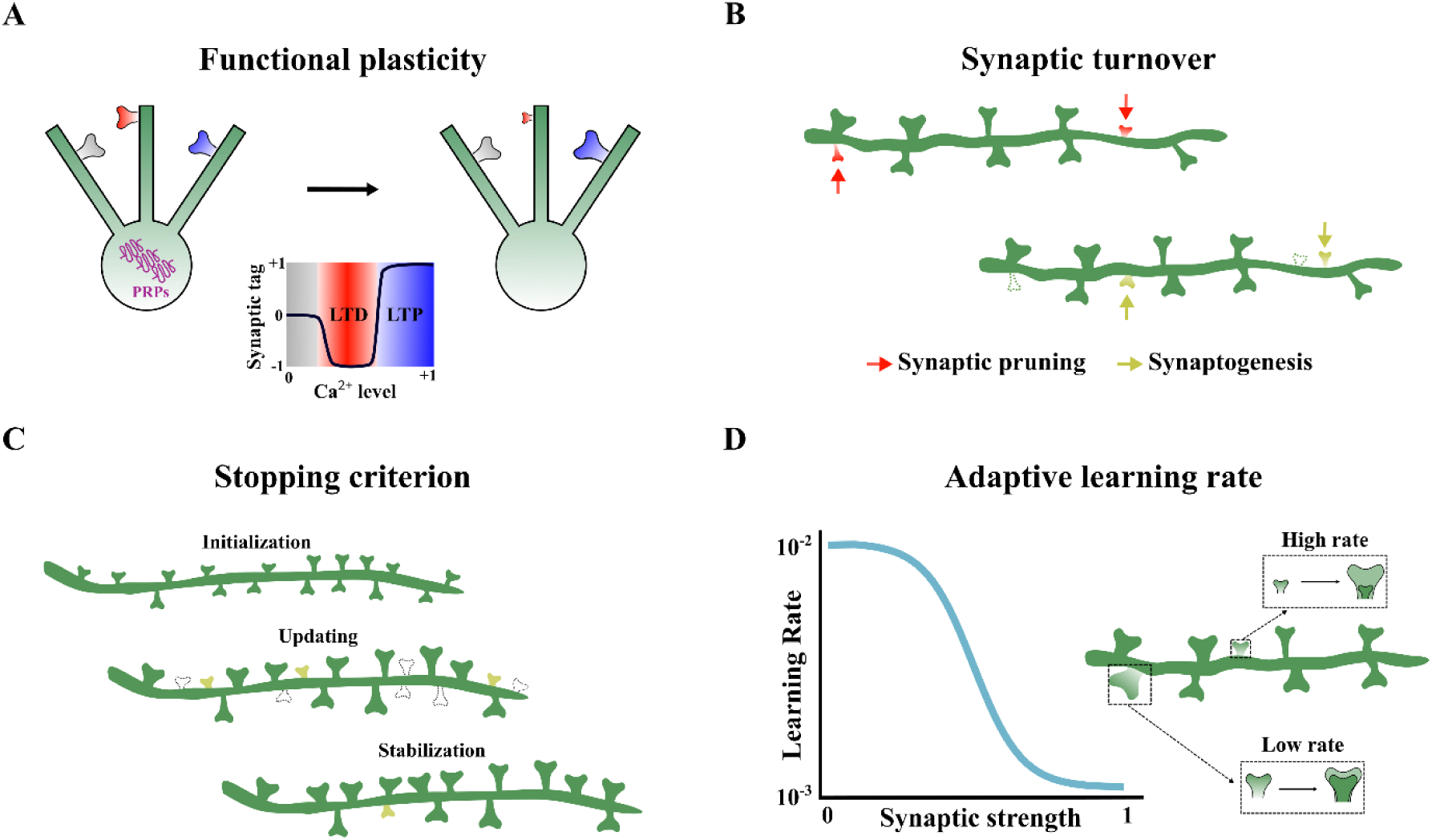
Functional and structural plasticity. **A.** Functional plasticity inspired by the synaptic tag-and-capture rule, as in^56^. **B.** Small synapses are systematically pruned and replaced with new ones at different random locations between the inputs and the pyramidal neurons’ dendrites. **C.** Bigger synapses are more stable as their learning rate decreases while their strength increases. **D.** The network is updated until half of the initial synapses become stable (their learning rate falls below the sigmoidal’s midpoint), where we assume stabilization of the circuit via learning and stop training.

### Synaptic turnover improves learning in a custom binary classification task

To explore the role of synaptic turnover in learning, we first trained and tested our network model on binary classification tasks of varying difficulty. We compared two conditions: one with synaptic turnover enabled and one in which turnover was disabled (**Figure 3**). The classification task consisted of a custom-based dataset of 20×20 pixel images depicting rectangles and circles of varying location and pixel intensity. The task difficulty was defined by the degree of overlap between rectangles and circles and the positional stability of the two classes (**Figure 3B**) (see Methods). Performance was estimated based on a majority rule, in the absence of a teaching signal, and using unseen images. Specifically, the predicted class label of a test image was determined by the subpopulation with the largest number of active neurons. A neuron was considered active if its firing rate was greater than 5Hz.

**Figure 3.**
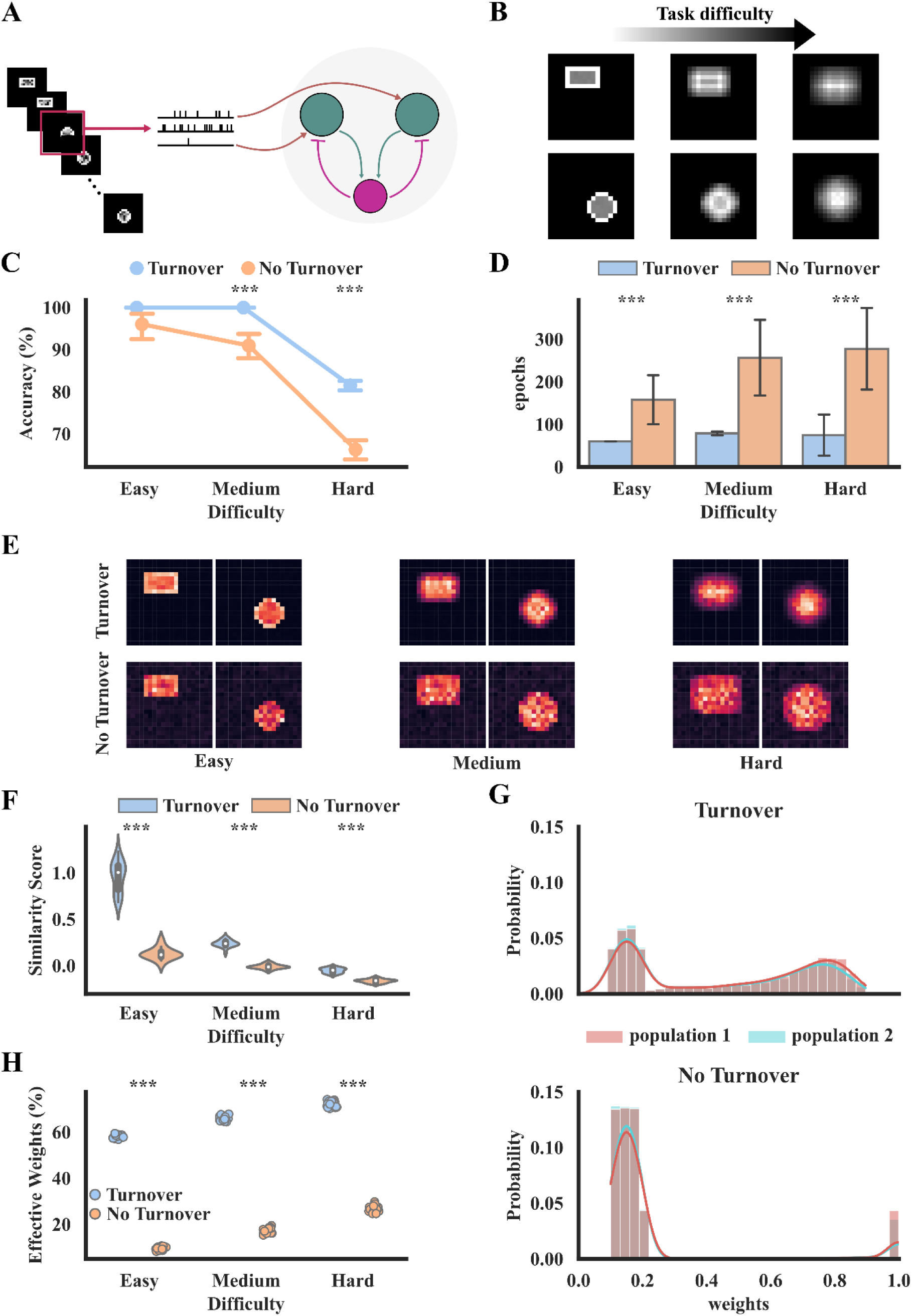
Synaptic turnover facilitates learning across various task difficulties. **A.** Schematic representation of the classification task. Images were translated into spike trains and fed into a network with or without synaptic turnover. **B.** Examples of the average image for each class as the difficulty level increases**. C.** Performance accuracy (on the test set) across three difficulty levels for both network models. Differences among groups are statistically significant (2-way ANOVA: turnover F(1, 114)=200.907, p<10^-3^, difficulty F(2, 114)=14.21, p<10^-3^, interaction F(2, 114)=7.871, p=10^-2^). Stars denote significance with unpaired t-test (two-tailed) with Bonferroni’s correction. **D.** Number of epochs (equal to the number of input images) to convergence. Differences among groups are statistically significant (2-way ANOVA: turnover F(1, 114)=115.6, p<10^-3^, difficulty F(2, 114)=306.43, p<10^-3^, interaction F(2, 114)=13.974, p<10^-3^). Stars denote significance with unpaired t-test (two-tailed) with Bonferroni’s correction. **E.** Representational maps of each subpopulation that correspond to the input class. Colormap denotes the summed synaptic weights corresponding to the specific input neuron (warmer colors denote larger weight values). **F.** Similarity score of the representational maps shown in **E.** The score is equal to the normalized difference between intra- and inter-cluster distance. Lower values denote a lower quality of the representational map (2-way ANOVA: turnover F(1, 114)=804.303, p<10^-3^, difficulty F(2, 114)=745.83, p<10^-3^, interaction F(2, 114)=246.267, p<10^-3^). Stars denote significance with unpaired t-test (two-tailed) with Bonferroni’s correction. **G.** Post-learning weight distributions for each subpopulation. **H.** Percentage of effective weights, namely those with values above a threshold level (>0.3) (2-way ANOVA: turnover F(1, 114)=60891.236, p<10^-3^, difficulty F(2, 114)=2192.982, p<10^-3^, interaction F(2, 114)=27.490, p<10^-3^). Stars denote significance with unpaired t-test (two-tailed) with Bonferroni’s correction.

We found that the presence of synaptic turnover significantly increased the classification accuracy, especially for higher levels of task difficulty (**Figure 3C**). In addition, synaptic turnover allowed faster learning across all difficulty settings, as shown by the number of training epochs (**Figure 3D**). Notably, our network is trained by presenting images one by one. Thus, the number of epochs equals the number of training samples. As a result, synaptic turnover not only improves performance accuracy and learning speed but also uses a much smaller training data set.

Having demonstrated that synaptic turnover facilitates learning as difficulty levels rise, we next sought to understand why this happens. Towards this goal, we generated representational maps for the two class-specific subpopulations of neurons across the three difficulty levels. These maps are generated by summing the weights of all input neurons, thus depicting the strength of the encoding for each pixel on the input space (see Methods). As such, they correspond to a 20×20 representation of what was learned by each subpopulation.

Visual inspection of the three difficulty cases suggests that the quality of representational maps is higher with synaptic turnover (**Figure 3E**). To quantify this improvement, we used a similarity metric, which compares the inter-class and the intra-class distances (see Methods). We found that synaptic turnover results in a higher similarity score - and thus better quality of representation - for all difficulty settings (**Figure 3F**). To assess whether this result is due to better utilization of trainable parameters, we looked at the weight distributions post-learning. Synaptic turnover results in a higher percentage of effective weights and a flatter distribution of post-learning weights (**Figure 3G-H**). These findings show that synaptic turnover enables better utilization of trainable weights, presumably because those that do not carry important information turnover until they become useful for the task at hand.

Taken together, these simulations suggest that synaptic turnover improves learning by optimizing the use of available resources (in this case, trainable weights), a feature that is particularly important for challenging tasks where each trainable parameter matters.

### Synaptic turnover is more beneficial for learning under conditions of resource scarcity

To gain a deeper understanding of how synaptic turnover may optimize the uses of available resources, we simulated conditions in which the number of available resources varies. Towards this goal, we increased (doubled) and decreased (halved) the number of trainable weights (control value: 1,000) in our SNN model, keeping all other parameters identical and repeating the experiments shown in **Figure 3**.

We found that when resources are halved, synaptic turnover results in major increases in performance accuracy across all difficulty levels. In fact, the SNN model is unable to solve the classification task without synaptic turnover under this condition, as its performance remains near chance across all difficulty settings (**Figure 4B** top, left). On the contrary, when resources are doubled, synaptic turnover does not provide any performance benefits (**Figure 4B** top). However, the beneficial effect of synaptic turnover on learning speed (and the number of training examples needed) is consistent across all resource and difficulty levels (**Figure 4B** bottom). Specifically, an SNN model with synaptic turnover can learn any task with as few as 100 training images. Without synaptic turnover, this number can increase by more than 3 times. This result aligns with theoretical observations that synaptic turnover is essential for learning and storing information, especially in sparsely connected networks like those found in the brain^57^.

**Figure 4.**
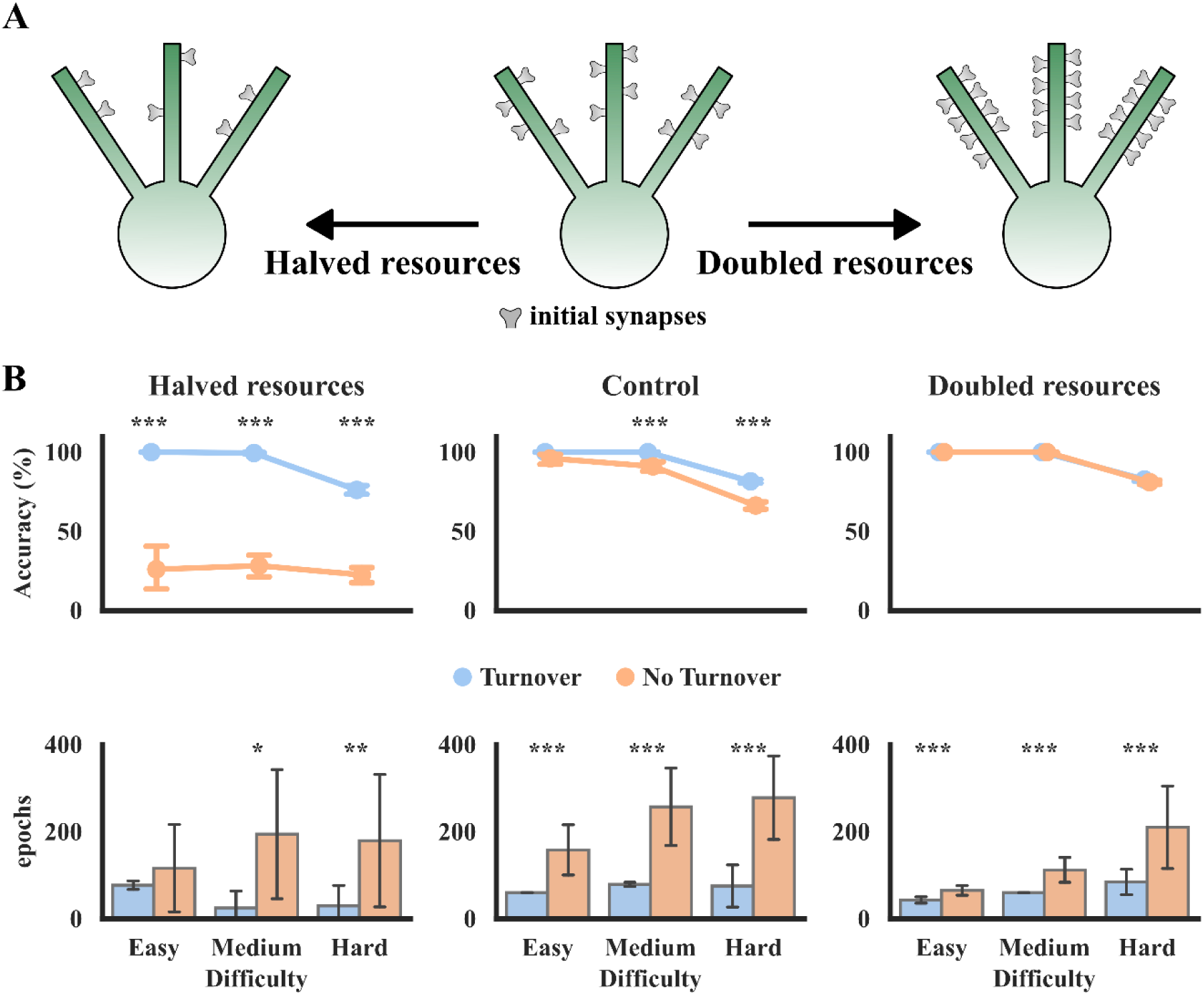
Synaptic turnover is more beneficial under conditions of resource scarcity. **A.** Schematic illustration of the SNN model with variable numbers of trainable weights (resources): halved (left), control (middle), and doubled (right). **B.** (top) Performance accuracy on the test set as a function of task difficulty for the three resource cases for SNN models with (blue) and without (orange) synaptic turnover (Halved: 2-way ANOVA: turnover F(1, 114)=561.831, p<10^-3^, difficulty F(2, 114)=11.358, p<10^-3^, interaction F(2, 114)=5.178, p=7·10^-3^; Control: 2-way ANOVA: turnover F(1, 114)=115.6, p<10^-3^, difficulty F(2, 114)=306.043, p<10^-3^, interaction F(2, 114)=13.974, p<10^-3^; Doubled: 2-way ANOVA: turnover F(1, 114)=3.701, p=5.7·10^-3^, difficulty F(2, 114)=1678.353, p<10^-3^, interaction F(2, 114)= 3.701, p=2.8·10^-3^). Stars denote significance with unpaired t-test (two-tailed) with Bonferroni’s correction. (bottom) Number of epochs (equal to the number of training images) to convergence for each task and resource case (Halved: 2-way ANOVA: turnover F(1, 114)=43.490, p<10^-3^, difficulty F(2, 114)=0.173, p=0.842, interaction F(2, 114)=5.016, p=8·10^-2^; Control: 2-way ANOVA: turnover F(1, 114)=200.907, p<10^-3^, difficulty F(2, 114)=14.21, p<10^-3^, interaction F(2, 114)=7.871, p=10^-3^; Doubled: 2-way ANOVA: turnover F(1, 114)=73.542, p=10^-3^, difficulty F(2, 114)=49.616, p<10^-3^, interaction F(2, 114)= 15.728, p<10^-3^).

Overall, the above simulations show that the beneficial effects of synaptic turnover on learning speed and the number of training examples are ubiquitous in our SNN configuration. Performance benefits, on the other hand, become especially pronounced when the available resources (i.e., plastic synapses) are limited. Given the high energy demands of plastic synapses^73^, our simulations suggest that synaptic turnover not only improves learning speed and performance but does so in an energy-efficient manner.

### Synaptic turnover improves learning in binary MNIST classification task

To further solidify our claims about the benefits of synaptic turnover in learning, we tested our network against a more challenging task using the MNIST dataset, a well-known machine-learning benchmark dataset consisting of handwritten digits^74^. In the results shown here, the model learns to discriminate between two classes, resulting in 45 binary classification tasks. As in the custom-made task, we test three SNN configurations using half the control and double the number of trainable weights (Control value: 1,750). We found that synaptic turnover improved the performance accuracy of the SNN classification for most digit pairs under two resource availability settings: the control case and the case when trainable parameters were halved (**Figure 5A**). Improvements are especially pronounced when resources are halved, in which case a model without synaptic turnover fails to classify most digit pairs correctly. This improvement in performance accuracy was also reflected in the robustness of learning across different initializations: synaptic turnover made the performance more stable in the control and halved-resources cases but not in the double-resources configuration (**Figure 5B**). These findings suggest that, as in our custom-made classification task, MNIST classification benefits from synaptic turnover primarily when the number of trainable parameters in the SNN model is small.

**Figure 5.**
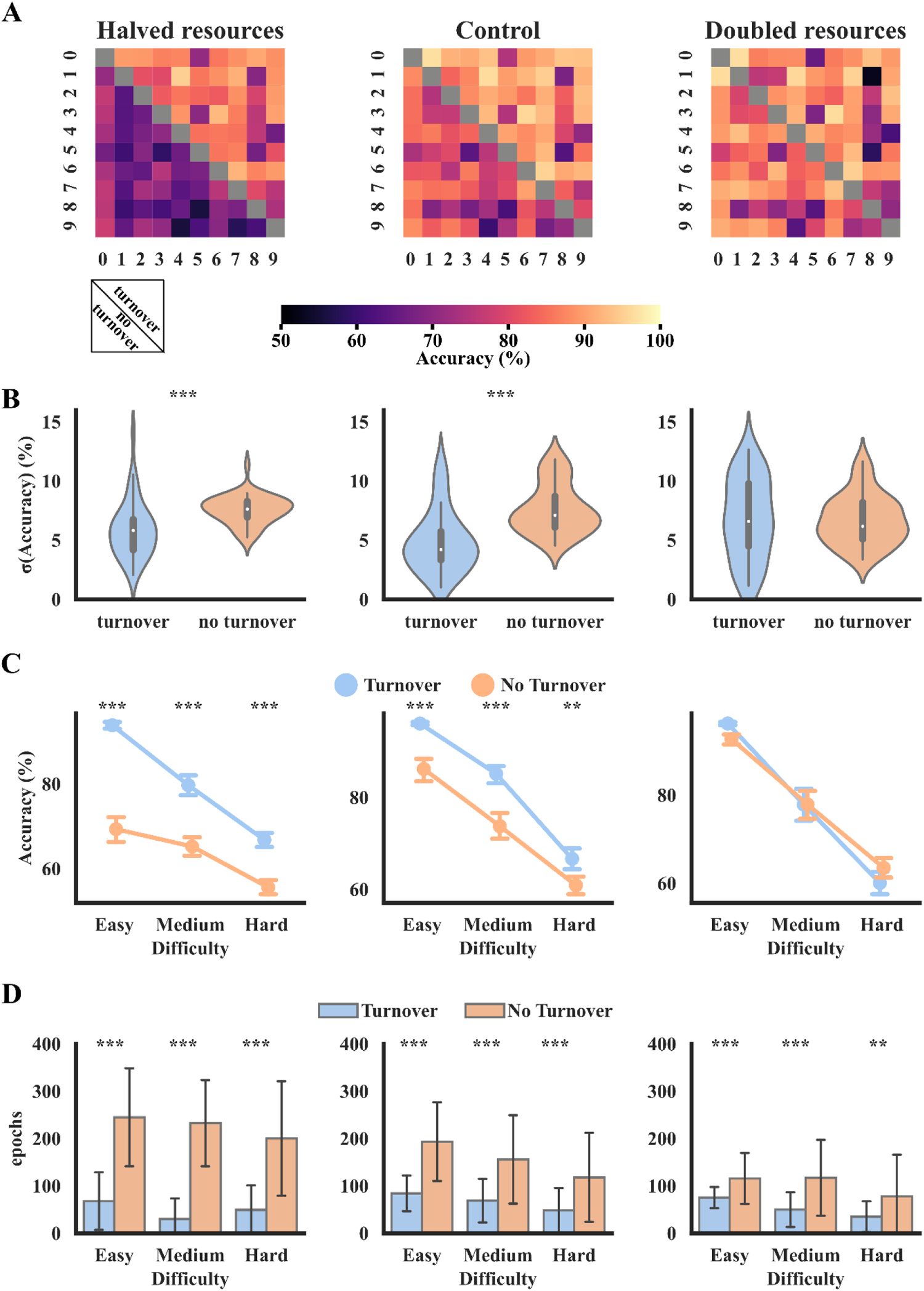
Synaptic turnover facilitates the learning of MNIST handwritten digits. For all sub-panels of this figure, the SNN model has half (left), control (middle) and double (right) the number of plastic synapses (trainable weights). **A.** Heatmaps of performance accuracy on the test. The upper triangle of the heatmap shows the performance with synaptic turnover, while the lower part shows the performance without synaptic turnover. **B.** Standard deviation of the accuracy across 20 random initializations of the model for all binary classification tasks (Halved: t-statistic(88)=4.281, p<10^-3^, p=0.911; Control: t-statistic(88)=5.544, p<10^-3^; Doubled: t-statistic(88)=0.112, p=0.911). **C.** Performance accuracy for three levels of difficulty: easy, medium, and hard across the three resource conditions. Easy (1 vs. 4, 3 vs. 6), medium (2 vs. 5, 3 vs. 8), and hard (5 vs. 8, 4 vs. 9) tasks are selected based on the average performance accuracy of the SNN on these tasks (Halved: 2-way ANOVA: turnover F(1, 234)=379.835, p<10^-3^, difficulty F(2, 234)=188.755, p<10^-3^, interaction F(2, 234)=21.867, p<10^-3^; Control: 2-way ANOVA: turnover F(1, 234)=104.796, p<10^-3^, difficulty F(2, 234)=323.709, p<10^-3^, interaction F(2, 234)=3.655, p=0.027; Doubled: 2-way ANOVA: turnover F(1, 234)=0.001, p=0.974, difficulty F(2, 234)=330.871, p<10^-3^, interaction F(2, 234)=3.778, p=0.024). **D.** Number of epochs to convergence across the different task difficulty levels and the three resource variability settings (Halved: 2-way ANOVA: turnover F(1, 234)=268.619, p<10^-3^, difficulty F(2, 234)=2.189, p=0.043, interaction F(2, 234)=1.905, p=0.151; Control: 2-way ANOVA: turnover F(1, 234)=268.619, p<10^-3^, difficulty F(2, 234)=3.189, p=0.043, interaction F(2, 114)=1.905, p=0.151; Doubled: 2-way ANOVA: turnover F(1, 234)=93.59, p<10^-3^, difficulty F(2, 234)=12.291, p<10^-3^, interaction F(2, 234)= 1.557, p=0.213).

To assess whether the synaptic turnover benefits extend to more challenging discrimination tasks, we clustered digits into three increasing-difficulty groups. Specifically, the *easy* task consisted of pairs 1 vs. 4 and 3 vs. 6, the *medium* task consisted of pairs 2 vs. 5 and 3 vs. 8, and the *hard* task consisted of pairs 5 vs. 8 and 4 vs. 9. We found that synaptic turnover resulted in higher performance accuracy for the control and halved resources configurations, but not when available resources were doubled (**Figure 5C**). However, as with our custom-made task, synaptic turnover increased the learning speed and reduced the number of training examples needed across all difficulty levels and resource availability configurations (**Figure 5D**).

To gain insights into how synaptic turnover facilitates learning, we constructed representational maps (averaging across all subpopulations that learned the same digit) for the control configuration. We found that synaptic turnover induced high-quality representational maps that were closer to their corresponding class, both by visual inspection (**Figure 6A**) and based on a similarity score (**Figure 6B**). As in our custom dataset case, improved quality of representational maps emerged from the better utilization of trainable weights endowed by synaptic turnover (**Figure 6C**). This was evident by the uniformity in the distribution of post-learning synaptic weights, while most of the trainable weights remained very small (unused) when synaptic turnover was disabled (**Figure 6D**).

**Figure 6.**
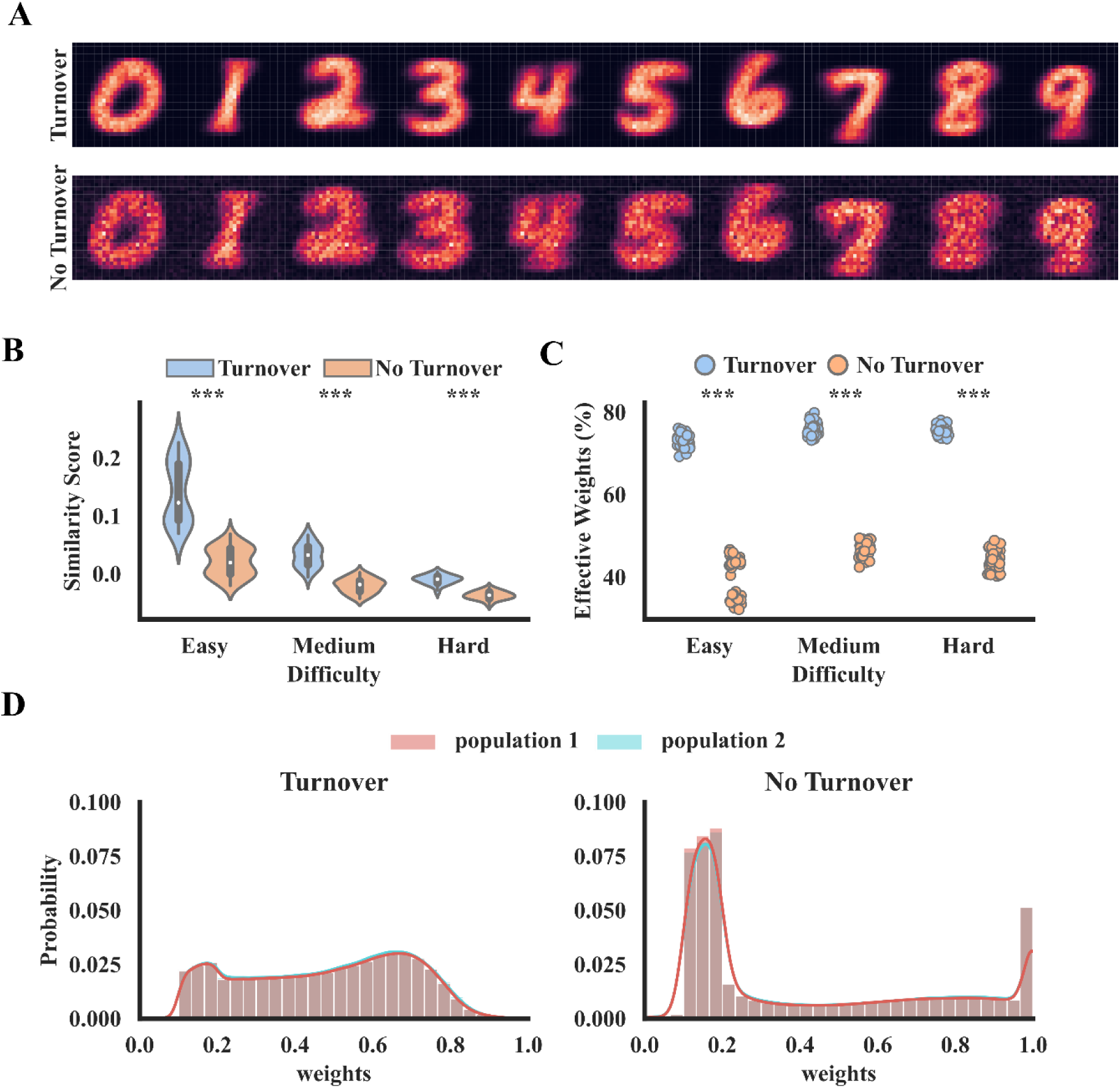
Synaptic turnover results in high-quality representations through better utilization of weights. **A.** Representational maps for the MNIST digits with (top) and without (bottom) synaptic turnover. Each digit participates in 9 pairwise comparisons, resulting in 9 representational maps. The average of these maps is shown here. **B.** Similarity score of the representational maps shown in **A.** The score is equal to the normalized difference between intra- and inter-cluster distance. Lower values denote a lower quality of the representational map (2-way ANOVA: turnover F(1, 234)=380.618, p<10^-3^, difficulty F(2, 234)=332.768, p<10^-3^, interaction F(2, 234)=67.017, p<10^-3^). **C.** Percentage of effective weights (>0.3) post-learning as a function of task difficulty level (2-way ANOVA: turnover F(1, 234)=8927.429, p<10^-3^, difficulty F(2, 234)=84.668, p<10^-3^, interaction F(2, 234)=15.119, p<10^-3^). **D.** Post-training weight distributions of the two SNN subpopulations. Distributions are generated across all tasks and initializations.

Taken together, our MNIST simulations show that the beneficial effects of synaptic turnover on the speed of learning and the number of training examples are consistent across all tasks and difficulty settings. However, performance accuracy and robustness benefits are pronounced under control and resource scarcity conditions but do not hold when the number of trainable parameters is high. A more in-depth analysis of the respective representational maps for each task revealed that synaptic turnover leads to higher-quality representations through the improved utilization of trainable weights. This is because synaptic turnover allows unused (small) weights to turnover until they convey information that is useful for the classification task, in which case they grow in size.

The above results highlight the importance of synaptic turnover in improving learning across a wide range of tasks and resource availability settings. However, certain MNIST digit pairs remained especially challenging even with synaptic turnover enabled in our SNN (**Figure 5A**). To find a solution to this problem, we turn to another feature of the biological brain that has not yet been incorporated into our SNN model, specifically that of distance-dependent axonal rewiring. A recent study has shown that neurons share the same activity pattern in temporal space and are organized as clusters in anatomical space^75^. Thus, it is reasonable to assume that an axon undergoing turnover will ultimately end up in a neighboring neuron. We modified the synaptic turnover mechanism to include a topology dependent constrained, i.e., constrained turnover. Specifically, a weak synapse belonging to a subpopulation will be replaced by another synapse in the same subpopulation. Evidence from such a constrained turnover comes from studies in which learning results in the formation of new synapses close to existing spines^76,77^, thus reinforcing the learning of the specific task. We found that the SNN model with constrained synaptic turnover classifies handwritten digits with higher accuracy than our random turnover model (**Figure 7A**), and this improvement is consistent across different initialization conditions (**Figure 7B**). When looking at especially challenging pairs, we see that constrained turnover improves performance by increasing both the accuracy and its robustness (**Figure 7C**), while requiring a few training examples (**Figure 7D**).

**Figure 7.**
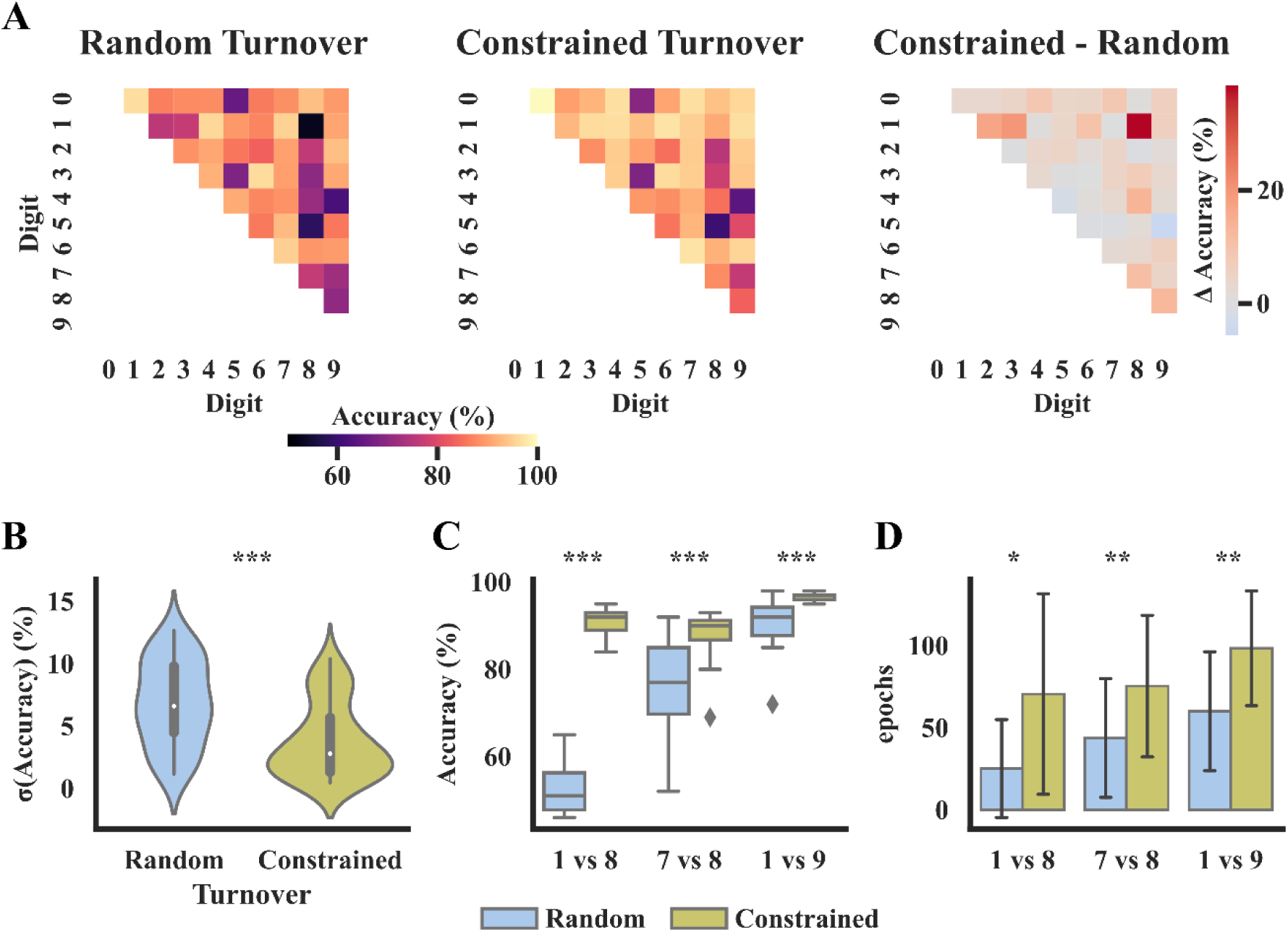
Constrained turnover results in higher accuracy and lower variability across almost all digit pairs. Available resources: 3,500 plastic synapses. **A.** Heatmaps of performance accuracy on the test set. (left) Performance with random synaptic turnover, (middle) with constrained turnover, (right) performance difference, Δ(constrained-random). **B.** Standard deviation of the accuracy across 20 random initializations of the model for all binary classification tasks (t-statistic(88)=4.407, p<10^-3^). **C.** Performance accuracy for three pairs that show improvements. In all presented cases the constrained turnover reduces the std of the performance accuracy (1 VS 8: t-statistic(38)=24.869, p<10^-3^; 7 VS 8: t-statistic(38)=3.97, p<10^-3^; 1 VS 9: t-statistic(38)=4.614, p<10^-3^). **D.** Number of epochs to convergence across the pairs shown in **C.**, for random and constrained turnover (1 VS 8: t-statistic(38)=2.987, p=0.005; 7 VS 8: t-statistic(38)=2.512, p=0.016; 1 VS 9: t-statistic(38)=3.403, p<10^-3^).

These results suggested that synaptic turnover is more efficient when synapse elimination and formation are restricted within its own subpopulation. This may explain why small-world networks are observed in many brain areas and across species^78^.

## Discussion

We investigated the impact of synaptic turnover, a type of structural plasticity, on the learning and classification performance of a bio-realistic SNN model with Ca^2+^-dependent learning rules and few compartment neurons. The SNN model was based on prior work^56,58^ and was extended to include a teaching signal, different interneuronal to pyramidal cell connectivity, and to receive image-based input signals. The model was applied to custom-based and MNIST binary classification tasks to understand how synaptic turnover may facilitate learning. While we focused on synaptic turnover, the proposed SNN model provides a useful framework for studying the role of various biological features in learning and classification. These include different learning rules, types of inhibition, wiring structure, compartmentalization, etc.

We found that the presence of synaptic turnover resulted in higher performance accuracy and faster learning across all classification tasks tested. It also allowed learning with a much smaller number of training examples to achieve the same performance as a network without synaptic turnover. Importantly, these learning improvements were highest under conditions of resource scarcity, namely when the number of trainable parameters was halved, and the task difficulty was increased. When trainable parameters were doubled, the contribution of synaptic turnover in performance accuracy diminished while speed and training sample benefits persisted. These improvements were due to a more accurate representation of the different image categories by the network weights through their flexible and more efficient utilization. Flexibility is enhanced because synapses that do not convey important information (due to random initialization) can turnover until they find an appropriate presynaptic partner, in which case they can effectively contribute to the task at hand. This process can be further facilitated by restricting turnover so that new synapses are formed within the sub-population in which they were eliminated, thus ensuring an even distribution of resources to both class-specific populations.

Recent advancements in neural networks incorporating bio-inspired learning rules and dendritic mechanisms have generated enthusiasm in the scientific community as they promise solutions to challenging problems like continual learning and credit assignment in the absence of error backpropagation^79–81^. However, the ways in which biological plasticity rules, anatomical and biophysical properties of the neurons cooperate to drive efficient learning remain explored. Towards this goal, we used a SNN model that incorporates various forms of functional and structural plasticity in multicompartmental neuronal models with active dendrites. We selected these because of their established impact on learning and memory processes^82,83^ and their prevalence across regions and species^84^. The proposed model can thus serve as a basis for a more extensive exploration of the integrative effects of these mechanisms in biological and machine learning.

Our study highlights the potential of using bio-inspired learning rules for machine learning applications. While currently, such training rules cannot match the performance of traditional algorithms for practical applications, this approach holds promise for future generations of SNN models. Our bio-inspired, supervised learning method requires a very small amount of labeled data, contrary to classical methods^85^, while achieving very high-performance accuracy. As such, it opens the way for the construction of complex SNN architectures that incorporate biological features like recurrent connections, cross-linked associations, reinforcement, attention, and other types of layers. Numerous studies have explored the development of local learning rules for SNNs^86–92^ (for recent reviews see^93,94^), as well as the hardware implementation of neuromorphic computing systems with analog weights and spiking architectures^95,96^ (for a review see^97^). Therefore, the development of SNN systems like the one proposed here, which are readily implementable on neuromorphic hardware devices, will facilitate the creation of real-time, energy-efficient information processing systems for various applications (e.g., communication, household appliances, industrial production, robotics)^98^.

It is crucial to acknowledge the limitations of our modeling approach. First, our model has only been evaluated on binary classification tasks. While increasing the size of our network in terms of neural units and synapses would allow us to tackle more complex tasks, such expansion is beyond the scope of this study, whose goal is to demonstrate the efficiency gains of synaptic turnover. Second, proving a teaching signal during training is not in line with single-area models, as teaching signals in the brain arise from already established networks, as demonstrated in various regions^99,100^. Ideally, these networks should be formed through unsupervised learning algorithms. A more biologically plausible alternative could be a self-supervised learning framework, where teaching signals arise from the similarity and dissimilarity of inputs rather than class labels directly^101^. Despite the limitations of our teaching signal approach, our study is still valuable as it allows us to evaluate the contribution of the synaptic turnover mechanisms to efficient learning. The demonstration that topologically constrained synaptic turnover can improve performance and facilitate learning has the potential to inspire the integration of such mechanisms into advanced architectures in the AI field.

The most important take-home message of this work is that synaptic turnover improves the efficiency of learning in our SNN model. This is important because brains evolved under evolutionary pressure and have limited resources (neurons, synapses, energy). Thus, coming up with mechanisms that allow accurate learning with the use of limited resources is critical for survival. Given the advent of technology, resource limitations may not apply to current machine learning systems. However, the fact that state-of-the-art ML systems, such as GPT-4, require the energy of a small city to be trained, charts a non-sustainable future for such AI models. The findings of this study can inspire future developments to adopt biological features to improve the efficiency and sustainability of ML systems.

## Methods

### Datasets

Our model was evaluated on two datasets. Initial experiments were conducted on a custom shape dataset that we had control over, and then, we solidified our claims by utilizing MNIST^74^, a well-known machine learning benchmark dataset.

#### Shape Dataset

The shape dataset consists of 20×20 pixel images that contain a rectangle at the top left denoting class 1 and a disk at the bottom right for class 2 (**Figure 3B**). The disks and the rectangles consist of bright pixels at maximum intensity on their perimeter and pixels with random intensity inside. The total amount of images generated per class was 1,000 for training and 200 for testing. The task’s difficulty (i.e., easy, medium, hard) was determined by two variables, the intraclass stability, and the interclass overlap. Stability corresponds to how much the mass center of a shape could move from image to image within the same class, negatively correlated with the task difficulty level, i.e., making the task easier as it increases since it makes the data per class less variable. Overlap, on the other hand, is a metric corresponding to the amount of potential overlap between the two classes, making the task harder as it increases since it increases the similarity between the two classes. In this work, we used three different combinations of stability and overlap to generate shape datasets. The easy task corresponds to overlap equals zero and stability equals one, the medium task to slight overlap and stability and the hard task to adequate overlap and low stability (**Figure 3B**, for the means per class depicted as images). For every simulation, a new shape dataset is generated to increase the robustness of our results.

#### MNIST handwritten digits

The MNIST (Modified National Institute of Standards and Technology database) is a well-known handwritten digit dataset widely used to benchmark machine learning models. It comprises 70,000 28×28 pixel images of handwritten digits from 0 to 9. The dataset is already split into a training set containing 60,000 images and a test set containing 10,000 images.

### Neuronal Model

The network consists of excitatory and inhibitory neuronal populations, modeled as two-stage integrators^8^, based on experimental data primarily from the hippocampal CA1 area. Neurons consist of a somatic spiking unit and independent dendritic subunits capable of nonlinear synaptic integration, dendritic spike initiation, and compartmentalized plasticity^60^.

Synaptic input in each dendritic subunit is integrated (Equation 1) and then passed to the somatic compartment.

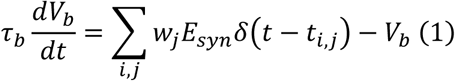

where *t_i,j_* denote the time *i* of incoming spikes in synapse *j*, *w*_*j*_ the weight of *j*-th synapse, *τ*_*b*_ the dendritic membrane time constant, *E_syn_* the unitary excitatory postsynaptic potential (EPSP), *V_b_* the *b*-th dendritic branch voltage, and *δ*(⋅) corresponds to Dirac delta function.

A depolarization component models the backpropagating action potential *V_b_^BP^*, which is added to all dendritic subunits connected to the somatic compartment. *V_b_^BP^* is modeled by an exponential decay (Equation 2),

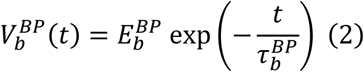

where *E_b_^BP^* is the peak of the backpropagating depolarization and *τ_b_^BP^* is the time constant of the backpropagating action potential.

The back-propagating action potential *V_b_^BP^* is summed with *V_b_* to determine the total depolarization of the dendrite, *V_d_*. When the sum of *V_b_* + *V_b_^BP^*, exceeds the dendritic spike generation threshold, *θ_dspike_*, a dendritic spike is generated, which causes the voltage of the corresponding subunit, *V_b_*, to rise instantaneously to *V_dspike_* (Equation 3).

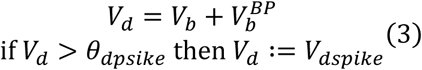

The voltage response of the somatic subunit and its spiking output is modeled by an integrate-and-fire point unit with adaptation^59^ (Equation 4).

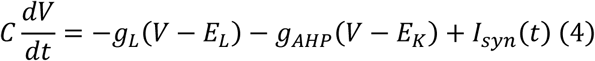

where *C* denotes the somatic membrane capacitance, *g_L_* the leak conductance, *E_L_* the somatic membrane resting potential, *g_AHP_* the conductance of the afterhyperpolarization (AHP) current, *E_K_* the potassium reversal potential, as AHP is a potassium-specific channel, and *I_syn_*(*t*) denotes the total synaptic current soma receives. Somatic spiking and the following reset occur when the somatic voltage crosses a threshold *V_T_*.

The adaptive conductance increases every time a somatic spike occurs (Equation 5).

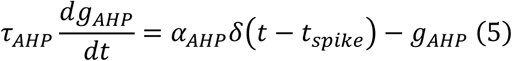

where *τ_AHP_* is the adaptation time constant, *α_AHP_* is the quantal increase of *g_AHP_* after a somatic spike occurs at time *t_spike_*. The time constant *τ_AHP_* can take two values corresponding to the neuron’s high and low excitability levels.

The synaptic input *I_syn_* to the somatic compartment is calculated based on the total dendritic synaptic current and the inhibition at the soma (Equation 6)

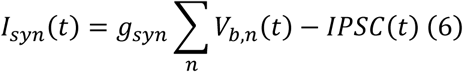

where *g_syn_* corresponds to the dendritic coupling constant, *V_b,n_* to the voltage of the *n*-th branch, and *IPSC* to the inhibitory input that the neuron receives.

### Synaptic Plasticity

#### Functional Plasticity

Learning in the model occurs according to the synaptic tag and capture plasticity model, which requires both synaptic tagging and the availability of protein-related proteins (PRPs) for stable strengthening and weakening of synapses^65^. When a postsynaptic branch receives presynaptic inputs, it leads to calcium influx. At the end of a pattern presentation event, a certain amount of calcium ions accumulated in the respective branches. Based on the individual calcium ion levels, a synaptic tag is generated, leading to long-term depression (LTD) of the corresponding synapse if the levels are low or long-term potentiation (LTP) if the levels are high. Once the synapses are tagged, plasticity-related proteins are produced that are consolidated together with the synaptic tags into the existing synaptic weights in the timespan of a couple of hours, leading to the synaptic weight update.

#### Calcium modeling

Calcium is critical to this plasticity mechanism, as it induces synaptic tags and PRP synthesis. The total calcium influx during the presentation of a pattern to a synapse determines the calcium Ca*_syn_*, which models the calcium concentration of each synapse. Each incoming synaptic spike causes a step increase of calcium, which depends nonlinearly on the local depolarization of the dendritic branch where the synapse resides. We assume that calcium influx upon arrival of a presynaptic spike ΔCa*_syn_* is primarily through NMDA receptors^102^, and is, thus, dependent sigmoidally on the dendritic membrane voltage (Equation 7).

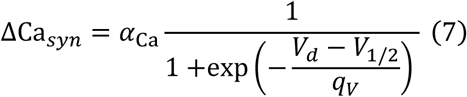

where *α_Ca_* denotes the maximum calcium ion influx, *V*_1/2_ and *q_V_* denote the midpoint and the steepness of the sigmoid function, and *V_d_* corresponds to the total dendritic membrane voltage. *V*_1/2_ = 30 mV and *q_V_* = 5mV are held constant for all simulations.

#### Plasticity Related Proteins Production

It has been shown in multiple plasticity studies that both dendritic^103,104^ and somatic^105,106^ protein synthesis is essential for synaptic weight consolidation, separately or even both at the same time^107^. The previously published model incorporated three different strategies for plasticity-related protein synthesis, where either somatic solely, dendritic solely, or both were taking place. In the current model, only the somatic PRP synthesis model was used. PRP synthesis initiation was modeled as an all-or-none phenomenon. When the calcium level of a dendritic branch, which is equal to the sum of all of its dendritic calcium levels, passes a threshold *P_soma_*, a PRP transient was generated as an alpha function (Equation 8).

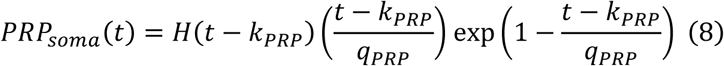

where *q_PRP_* denotes the steepness of the function, *K_PRP_* the shift in time, i.e., delay, and *H*(⋅) denotes the Heaviside step function. *K_PRP_* = 20 minutes and *q_PRP_* = 30 minutes for all simulations in the study.

#### Synaptic Tag Generation and Consolidation

As previously mentioned, the synaptic tag’s strength and sign are determined according to the Calcium Control Model^108^, thus lower levels of Ca^2+^ cause LTD, while higher levels cause LTP (Equation 9). The synaptic tag does not immediately alter the synapse’s weight but only after capturing PRPs, which are required for consolidation^109^.

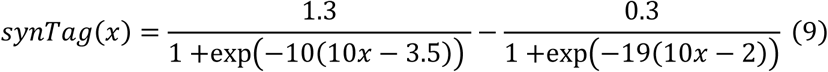

where *x* denotes the Ca^2+^ concentration.

Synaptic tags in the model decay exponentially with a time-constant *τ_synTag_* = 1 hour (Equation 10).

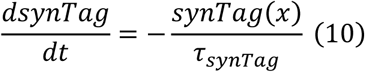

The synaptic tags are consolidated into synaptic weights over time (Equation 11).

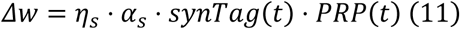

where *η_s_* is the learning rate of the synapse, *α_s_* the rate of synaptic tag consolidation, *synTag* the value of the synaptic tag at time *t*, either be positive or negative, and *PRP* to the PRP levels at time *t*. The weights are clipped in the range [0,1].

#### Dynamic Excitability

Learning has been shown to increase the excitability of neurons participating in forming a given memory^110–115^. On the other hand, neurons with increased excitability are more likely to participate in the formation of a new memory engram^112,116,117^. The activation of the transcription factor CREB (cAMP Response Element-Binding Protein) has also been found to modulate the excitability of neurons^118,119^ through the reduction of the AHP current^112,117^. Therefore, it has been suggested that learning makes cells more amenable to be recruited in future learning events by activating CREB^114,120–123^. Finally, it has been proposed that CREB may also induce the downstream activation of its own repressors^112,114^, which would reduce excitability after a certain period, thus creating a time window of increased neuronal activity excitability. Increased excitability is simulated through the transient reduction of the AHP current in the neurons in which PRP synthesis is triggered for approximately 12 hours after a learning event^56^.

#### Local Adaptive Learning Rate

A feature of biological synapses is that as they grow and become more stable, their growth rate drops^68,124–127^. This is incorporated into the network in the form of an adaptive and synapse-specific learning rate that is dependent on the synaptic weight value *w_s_*· *η_s_*(*w_s_*) corresponds to the learning rate of synapse s with weight *w_s_* and follows a reversed sigmoidal function (Equation 12).

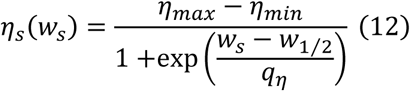

where *η_min_* and *η_max_* are the minimum and maximum learning rate values, respectively, *q_η_* and *w*_1/2_ denote the steepness and midpoint of the reversed sigmoidal, respectively.

#### Biologically Inspired Stopping Criterion

Larger spines have been observed to be more stable than smaller ones, leading to the hypothesis that the large ones should play a role in memory formation^33,68,124,126^. Given that, a stopping criterion was implemented that terminates training when a percentage of synaptic spines becomes larger than a threshold, assuming that memory has already been formed under such circumstances. A synapse *s* is characterized by its weight *w_s_* or its learning rate *η_s_*.

### Structural Plasticity

#### Synaptic Turnover

Synaptic turnover is a structural plasticity mechanism that involves synaptic pruning and synaptogenesis. After a given number of iterations *N_turn_*, plastic synapses with a weight smaller than the threshold *θ_turn_*, are pruned. Then, synaptogenesis takes place, where synapses equal to the number of pruned synapses are born between random presynaptic input neurons and postsynaptic neuron dendrites, with weight values pulled from a uniform distribution in the range [*w_min_*, *w_max_*].

### The Network

#### Network Model and Structure

The network consists of one input layer and a hidden/output layer. The input layer is connected to the hidden one sparsely, and learning occurs with the help of a class-specific learning signal during training. The learning signal is silenced during the testing phase, and the output evaluation occurs based on a majority rule.

#### Input Translation and Transmission

To use the datasets mentioned above as inputs for our spiking neural network, they are first flattened into a 1D vector. Then, they have to be translated into spike trains. This was done by translating pixel intensity into firing rates (Equation 13) and then generating a periodic spike train per pixel with the corresponding firing frequency. The stimulus duration was set to 4 seconds.

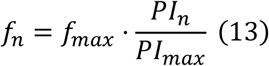

where *f_n_* is the firing rate of the n-th input neuron, and *PI_n_* denotes the pixel intensity of the *n*-th pixel.

The translated input is then transmitted to the hidden layer in spikes through synaptic connections. After the pattern has been presented, a consolidation period of 138 minutes follows. Synaptic tags and plasticity-related proteins are produced and consolidated into synaptic weights, leading to weight updates. After this period, the weights are updated, and the next input pattern is presented.

#### Network Connectivity and Computations

The network consists of an input and a hidden layer. The hidden layer possesses both excitatory pyramidal neurons and inhibitory interneurons. Input neurons are sparsely and randomly connected to the dendrites of the pyramidal neurons of the hidden layer, with initial weights pulled from a uniform distribution in the range [*w_min_*, *w_max_*] and those are the only plastic weights of the network.

For learning to occur, the network is equipped with a supervised mechanism as a teaching signal specific to each class. Each teaching signal targets a distinct subpopulation of pyramidal neurons in the hidden layer and fires with a firing rate *f_tn_*. This ensures that those neurons in the hidden layer that receive inputs both from the input layer and the learning signal will participate in the learning procedure of the presented pattern, leading to the formation of a memory engram. In this project, each task is a binary classification task. Thus, the teaching signal neurons and the corresponding pyramidal subpopulations were two.

Besides pyramidal neurons, interneurons are also represented in the hidden layer. The network possesses both dendritic and somatic targeting interneurons. Dendritic targeting interneurons have supralinear dendrites, while somatic targeting have sublinear dendrites.

In addition to the interneuron classification presented above, interneurons are also split into two functional groups. One group suppresses the network’s activity, as the network itself is quite excitable, and the other performs feedback inhibition to further assist with separating the two classes. The former consists of both dendritic and somatic targeting interneurons that randomly receive input from pyramidal neurons in the hidden layer, and, in return, they randomly project back to them. The latter one consists of exclusively dendritic targeting interneurons that are grouped in the same manner as pyramidal subpopulations are, while they receive input from one of the subpopulations each and inhibit the other one.

#### Network Training and Testing

The network’s training procedure has already been described more or less. Patterns are presented for 4 seconds each, with consolidation periods in between, until enough weights grow large enough for the stopping criterion to kick in and to stop the training procedure.

After training is finished, the model is evaluated in its ability to classify the patterns that it was presented with into its respective classes. To do that, the plastic weights are frozen, and the hidden layer receives input only from the input layer. For every testing pattern presented, the network’s decision is determined by a majority rule that compares the number of active neurons in each subpopulation and selects the digit corresponding to the subpopulation with the most active ones. An active neuron in our framework corresponds to a pyramidal neuron that fires with a firing rate higher than 5 Hz.

### Analysis and Visualization Techniques

#### Representation Maps

We generated the representation maps to visualize the representations formed in each subpopulation during learning. To generate them, we used the weights of the synapses projecting to each subpopulation. Specifically, for each input node, we summed the outgoing weights towards each subpopulation separately. Then, we rearranged the sums of the weights into 2D matrices of the same size as the original input images, resulting in two different representation maps per run, one for every subpopulation.

#### Similarity score

To quantify the quality of our representation maps, we defined a similarity score inspired by the silhouette coefficient used to quantify clustering quality^128^. The similarity score was calculated as a function of the within-class and the across-class distance metrics (Equation 14).

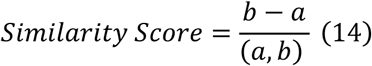

where *a* and *b* denote the average within-class and across-class distances, respectively.

Firstly, we measure the normalized Euclidean distance of both representation maps to every image in the test set. Then, we calculated the average distances 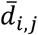 of the representation maps from the test images they encoded and test images belong to the opposite class (Equation 15).

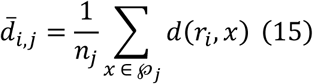

where *d*(*r_i_, x*) denotes the Euclidian distance between the representation map of class *i* and a test image *x* belongs to class *j*, and *n_j_* is the total number of test images of class *j*. *℘_j_* denotes the class *j*.

The within-class and the across-class distances are calculated as averages across classes, i.e., two classes in the case of the binary classification task (Equations 16).

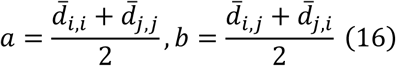

Where 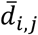 is the average Euclidean distance of the representation map of subpopulation *i* to every image in the test set belonging to class *j*.

### Computational Resources

The simulations for this project were run on the high-performance computing (HPC) cluster^129^ (version 7.0, http://www.rocksclusters.org/) of the Poirazi Lab, which consists of 624 cores and 3.25 TB shared RAM operating under CentOS Linux distribution. Data analysis was performed in Python (version 3.9) using the NumPy^130^, SciPy ^131^, and Pandas^132^ libraries. The plots were generated using Seaborn^133^ and Matplotlib^134^ libraries, and the statistical analysis was performed using Pingouin library^135^.

### Statistical Analysis

For all standard statistical tests (detailed in figure legends), the significance level *α* was 0.05. To correct for multiple comparisons, *α* was divided by the number of tests according to the Bonferroni procedure. Throughout the figures, p values are denoted by * (p<0.05), ** (p<0.01), and *** (p< 0.001). To compare performance among different groups (difficulty level x synaptic turnover), accuracies were analyzed using two-way analysis of variance (ANOVA) followed by an unpaired t-test (two-tailed) with Bonferroni’s correction whenever statistical difference was observed for posthoc comparisons.

## Constant and Parameter Tables

**Table 1.** Neuronal models parameters.

| Parameter | Description | Value |
| --- | --- | --- |
| $\tau_b$ | Passive dendritic integration time constant | 20 ms |
| $E_{syn}^{pyr}$ | Maximum unitary PSP of pyramidal cells | 4 mV |
| $E_{syn}^{int}$ | Maximum unitary PSP of interneurons | 3 mV |
| $\theta_{spike}^d$ | Voltage threshold for dendritic spike | 25 mV |
| $V_{spike}^d$ | Dendritic spike max depolarization | 50 mV |
| $E_L$ | Somatic leakage reversal potential | 0 mV |
| $\theta_{spike}^s$ | Voltage threshold for somatic spike | 20 mV |
| $g_{syn}^{pyr}$ | Dendritic coupling conductance of pyramidal cells | 180 nS |
| $g_{syn}^{int}$ | Dendritic coupling conductance of interneurons | 180 nS |
| $\tau_{soma}$ | Somatic time constant | 120 nS |
| $\tau_{AHP}$ | Adaptation time constant | 110-180 ms (fast, slow) |
| $\alpha_{AHP}$ | Spike-triggered adaptation conductance | 0.18 nS |
| $E_K$ | Adaptation reversal potential | -10 mV |
| $\tau_{bAP}$ | Back propagating action potential time constant | 17 ms |
| $E_{bAP}$ | Back propagating action potential max amplitude | 30 mV |

**Table 2.** Functional and structural plasticity parameters.

| Parameter | Description | Value |  |
| --- | --- | --- | --- |
| $\alpha_{Ca}$ | Calcium influx rate | 1.1 ms<sup>-1</sup> | |
| $P_{soma}$ | Threshold for somatic PRP synthesis | 18 a.u. | |
| $\tau_{PRP}$ | Time constant for PRP decay | 60 mins | |
| $\alpha_s$ | Rate of synaptic tag consolidation | 6.7 mins | |
| $\tau_{CREB}$ | Duration of increased excitability | 12.5 hours | |
| $[w_{min}, w_{max}]$ | Minimum and maximum initial weights for plastic synapses | [0.1, 0.2] a.u. | |
| $[\eta_{min}, \eta_{max}]$ | Minimum and maximum learning rate | [0.001, 0.01] a.u. | |
| $q_\eta$ | Slope of learning rate reverse sigmoidal | 10 a.u. | |
| $w_{1/2}$ | Midpoint of learning rate reverse sigmoidal | 0.5 a.u. | |
| $N_{iters}^{turn}$ | Number of iterations to synaptic turnover event | Shapes | MNIST |
|  |  | 10 | 20 |
| $w_{turn}$ | Weight threshold value for synaptic turnover candidate | 0.2 a.u. | |
| $P_{stop}$ | Percentage of large synapses required for stopping criterion | 50% | |
| $\eta_{large}$ | Learning rate threshold for a synapse to be considered as large | 0.0055 a.u. | |

**Table 3.**
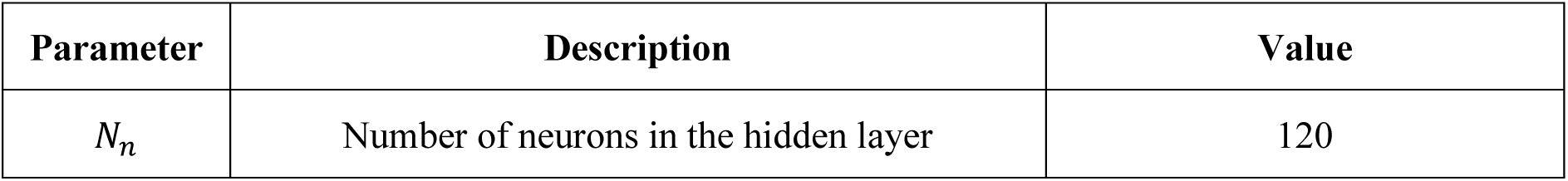
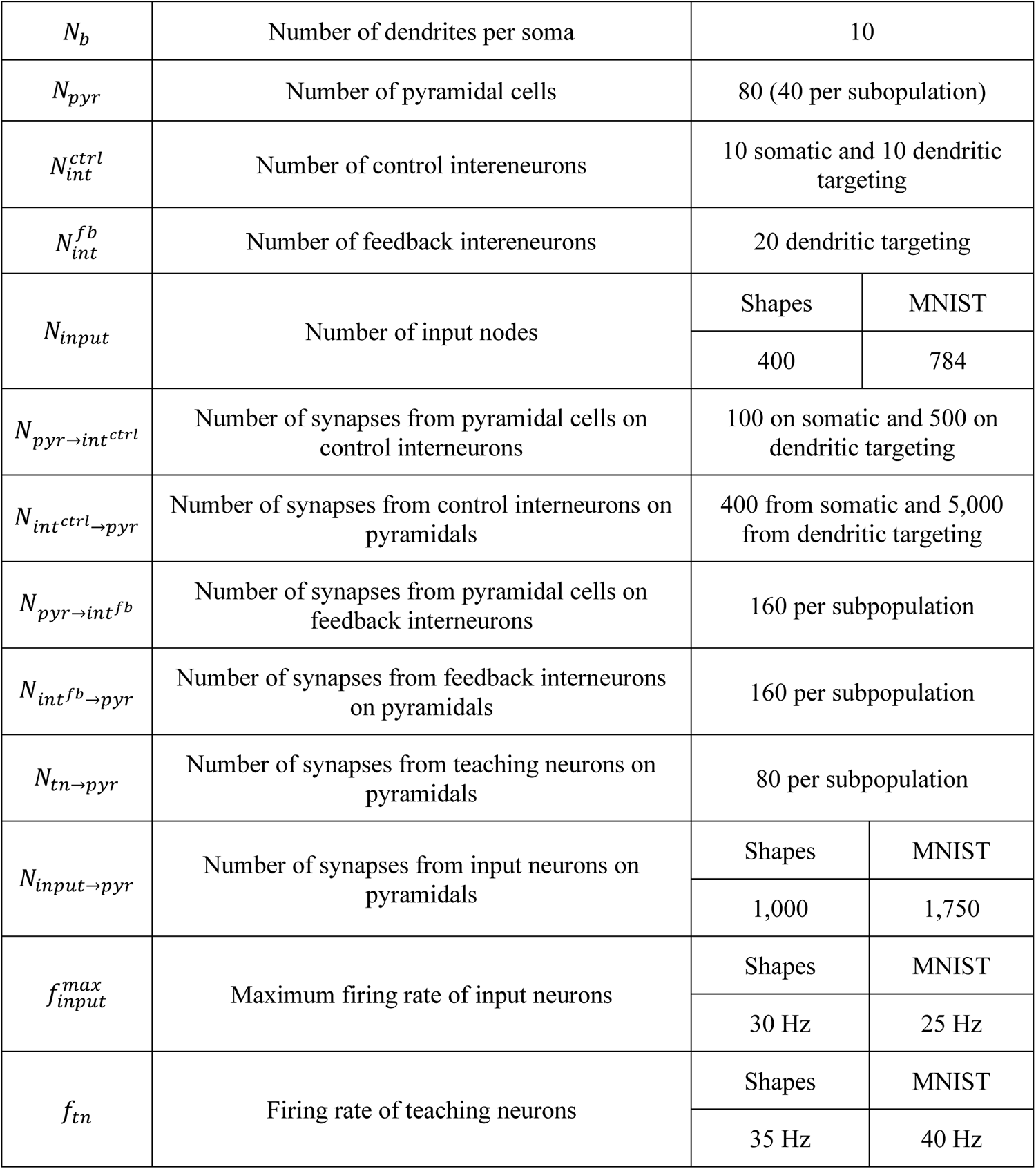
Network parameters.

| Parameter | Description | Value |
| --- | --- | --- |
| $N_n$ | Number of neurons in the hidden layer | 120 |

|  |  |  |  |
| --- | --- | --- | --- |
| $N_b$ | Number of dendrites per soma | 10 | |
| $N_{pyr}$ | Number of pyramidal cells | 80 (40 per subpopulation) | |
| $N_{int}^{ctrl}$ | Number of control interneurons | 10 somatic and 10 dendritic targeting | |
| $N_{int}^{fb}$ | Number of feedback interneurons | 20 dendritic targeting | |
| $N_{input}$ | Number of input nodes | Shapes | MNIST |
|  |  | 400 | 784 |
| $N_{pyr \rightarrow int^{ctrl}}$ | Number of synapses from pyramidal cells on control interneurons | 100 on somatic and 500 on dendritic targeting | |
| $N_{int^{ctrl} \rightarrow pyr}$ | Number of synapses from control interneurons on pyramidal cells | 400 from somatic and 5,000 from dendritic targeting | |
| $N_{pyr \rightarrow int^{fb}}$ | Number of synapses from pyramidal cells on feedback interneurons | 160 per subpopulation | |
| $N_{int^{fb} \rightarrow pyr}$ | Number of synapses from feedback interneurons on pyramidal cells | 160 per subpopulation | |
| $N_{tn \rightarrow pyr}$ | Number of synapses from teaching neurons on pyramidal cells | 80 per subpopulation | |
| $N_{input \rightarrow pyr}$ | Number of synapses from input neurons on pyramidal cells | Shapes | MNIST |
|  |  | 1,000 | 1,750 |
| $f_{input}^{max}$ | Maximum firing rate of input neurons | Shapes | MNIST |
|  |  | 30 Hz | 25 Hz |
| $f_{tn}$ | Firing rate of teaching neurons | Shapes | MNIST |
|  |  | 35 Hz | 40 Hz |

## Data and Code Availability

The source code that generates all Figures and the data supporting this study will be accessible on GitHub upon publication.

## Acknowledgments

This work was supported by NIH (1R01MH124867-02) to P.P., the European Commission (H2020-FETOPEN-2018-2019-2020-01, FET-Open Challenging Current Thinking, NEUREKA GA-863245) to P.P., and the Einstein Foundation Berlin, Germany (visiting fellowship, EVF-2019-508) to P.P.

## Author contributions

N.M., S.C., and P.P. conceived the project and designed the experiments. N.M. developed the model, performed the simulated experiments, and analyzed the data. N.M. and S.C. designed the figures and wrote the initial draft of the paper. S.C. supervised N.M. P.P. edited the manuscript, supervised and funded the project. All authors contributed to the article and approved the submitted version.

